# HALPred-B: Host-Aware Linear B-Cell Epitope Prediction: Challenges, Limitations, and Variability Across Species

**DOI:** 10.64898/2026.06.22.733770

**Authors:** Purnima Gautam, Ishaan Sinha, Pralay Mitra

## Abstract

Predicting linear B-cell epitopes is a basic immunoinformatics task that has a direct impact on vaccine design and antibody engineering. Recent advances in machine learning have improved predictive performance, but most existing approaches are trained on aggregated datasets and assume that antigenic patterns are conserved across host organisms. This assumption ignores the immunological variability depending on the host and prevents generalizing the model across species.

This is the first systematic host-wise evaluation where we present a systematic machine learning-based analysis of host-aware linear B-cell epitope prediction using curated datasets from the Immune Epitope Database (IEDB). We build separate datasets for human, mouse, and non-human primate hosts and assess several classification models, including Random Forest, Support Vector Machine (SVM), Gradient Boosting, XGBoost, and K-Nearest Neighbors (KNN). The models exploit feature representations derived from sequences, such as AAIndex descriptors, biochemical properties from ExPASy, and dipeptide composition.

Our results show that predictive performance differs substantially across hosts. Models achieve up to 86.07% accuracy and 0.93 ROC-AUC on human datasets but lower performance on mouse and non-human primate datasets. This gap underlies dataset bias and sequence distribution differences, as well as the inability of existing features to capture host-specific immunological context.

These results indicate that the prediction of linear B-cell epitopes is intrinsically host-specific, and a single global model does not generalize well across species. We propose to incorporate host-aware modeling strategies and organism-specific features for enhanced predictive reliability and biological relevance.

## I. Introduction

Linear B-cell epitopes are defined as contiguous stretches of amino acids that can be directly bound by antibodies and consequently are fundamental components of adaptive immunity. Their structural simplicity, which does not require immunogenicity of the native protein fold, makes them attractive targets for peptide-based vaccine design, immunodiagnostics, and therapeutic antibody development [1], [2]. These epitopes have traditionally been identified by laborious experimental techniques, including ELISA, peptide microarrays, and X-ray crystallography. These methods give high-resolution data but are costly, slow, and not suitable for proteome-scale applications [3], [4].

Hence computational prediction methods have become indispensable. In the past two decades, the field has progressed from physicochemical propensity scales to sophisticated machine learning frameworks. Early approaches, such as the Kolaskar-Tongaonkar algorithm, Parker hydrophilicity scale, and residue-level flexibility scores [5–8], were biologically interpretable but only marginally better than random, especially on large, diverse benchmarks [9]. Later machine learning models, such as BCPred [10], BepiPred [11], LBtope [12] and SVMTriP [15] showed better results via more complex feature representations such as amino acid compositions, dipeptide frequencies, and kernel-based string representations. More recently, taxonomy-aware approaches like epitope1D [16], phylogeny-informed predictors [17], organism-specific models [18], host-specific Nodaviridae predictors [19], and structurally enriched frameworks like BCEPS [20] have pushed the performance even further.

But an underlying assumption persists in almost all methods published to date: that patterns of antigens are conserved throughout the host organism in which the immune response takes place. This is biologically unreasonable to assume. Immune repertoires, antibody diversity, antigen-presentation pathways, and post-translational environments differ significantly among humans, mouse and non-human primates (NHPs) [21]–[23]. A peptide that induces a strong antibody response in a mouse may be immunologically silent in a primate and vice versa. When training data are pooled across hosts, as they almost always are, the resulting model inherits the distributional bias of the dominant organism, typically humans, and generalizes poorly to others.

This paper directly addresses this limitation. We make three specific contributions:

- We construct host-stratified benchmark datasets for human, mouse, and NHP hosts from IEDB and compare five standard classifiers under identical conditions, providing the first systematic, host-separated performance comparison in the literature.
- We quantify the cross-host accuracy gap and attribute it to differences in dataset size, shifts in sequence distribution, and lack of host-specific context in existing feature representations.
- We present Host-Aware Feature Reweighting (HAFR), a lightweight post-training approach that rescales feature vectors by host-derived importance scores to recover 2-4 percentage points of accuracy without retraining from scratch.

Taken together, these results support a clear conclusion: linear B-cell epitope prediction has to be considered as a host-dependent problem and not as a universal problem.

## II. Related Work

### A. Classical and Propensity-Scale Methods

The first linear B-cell epitope predictors were mostly based on residue-level physicochemical properties. The Hopp-Woods hydrophilicity scale [8] and Parker hydrophilicity method [5] used sliding windows to find hydrophilic regions that were likely to be exposed on the protein surface. Karplus and Schulz [6] used local flexibility as a proxy for accessibility and Kolaskar and Tongaonkar [7] proposed a semi-empirical antigenicity scale combining several physicochemical signals. Their simplicity and interpretability makes them still attractive, but large-scale benchmarking demonstrated that their performance is often only marginally better than chance on diverse datasets [9].

### B. Machine Learning Approaches

Supervised learning clearly outperformed classical propensity-based methods. BCPred [10] and BepiPred [11] showed that hidden Markov models and SVM-based string kernels can perform better when trained over enough labeled data. LBtope [12] used SVM classifiers with different feature sets and showed that fixed-length peptide representations with compositional features generalized more reliably than variable-length inputs. SVMTriP [15] enhanced specificity by combining tri-peptide antigenicity scores and similarity-based propensity features. ABCPred [14] employed recurrent neural networks for capturing short-range dependencies of epitope sequences and achieved competitive results on the benchmark datasets available at that time. However, these models were mostly trained on pooled datasets from multiple organisms, making their performance less clear in host-specific settings.

### C. Organism-Specific and Taxonomy-Aware Predictors

Recent studies are starting to incorporate biological context more directly. The epitope1D framework [16] exploits taxonomic ontology and graph-based pathogen signatures to improve generalization across taxonomic groups. Campelo et al. [17] showed that phylogenetic relationships can help to predict orthopoxvirus epitopes by identifying common antigenic patterns between related pathogens. Ashford et al. [18] have shown that organism-specific models (e.g., separate models for viral and bacterial antigens) tend to perform better than universal models. Shih et al. [19] further applied this concept to host specificity in Nodaviridae by using host-derived traits from insects, fish and prawns. BCEPS [20] also adds structural and immunological features, including the likelihood of MHC-II presentation and glycosylation propensity, aiming to improve predictions for specific biological contexts.

### D. Problem Solved by This Work

These studies do acknowledge biological context but are mainly focused on pathogen taxonomy or grouping at the organism level. They do not systematically test whether machine learning models trained on one host generalize to another with host-separated training and test sets. Most evaluations pool human, mouse, and other host data, masking host-specific behavior in aggregate accuracy. This work puts that gap directly to bed.

## III. Problem Formulation and Motivation

Let 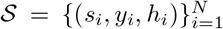 a collection of labeled peptide sequences be denoted by where *s*_*i*_ is a peptide sequence, *y*_*i*_ ∈ {0, 1} is the binary epitope label, and *h*_*i*_ ∈ ℋ = {human, mouse, NHP} specifies the host organism. Standard epitope predictors learn a single classifier *f*: *S* → {0, 1}by marginalising over *h*, implicitly treating host identity as irrelevant. We argue this is problematic for four distinct reasons:

1. **Host bias in public repositories**. IEDB [24] contains a heavily disproportionate number of human-derived assay entries. A model trained on the full IEDB implicitly optimizes for human immunogenicity, embedding human-centric biases that are invisible in aggregate metrics.
2. **Sequence distribution shift across hosts**. The distributions of amino acid composition and dipeptide frequencies differ measurably across host-derived epitope sets due to differences in antigen origin, assay selection, and immune selection pressure. A model trained on one distribution will not necessarily generalize to another.
3. **Absence of host-specific feature signal**. Physicochemical and compositional features encode intrinsic peptide properties but provide no information about the antibody repertoire, B-cell receptor diversity, or processing environment of the host. The model therefore cannot distinguish immunogenically relevant from irrelevant residues in a host-dependent manner.
4. **Overgeneralization of global models**. Across-species pooling conflates distinct immunogenic patterns, forcing the model to learn a compromise decision boundary that is suboptimal for every individual host.

The central question we investigate is therefore: *how does linear B-cell epitope prediction performance vary across host organisms under controlled, host-separated evaluation, and what are the limitations of current sequence-based feature representations in capturing host-specific immunogenic patterns?*

## IV. Methodology

Figure 1 provides an overview of the HALPred-B pipeline. Each stage is described below.

**Fig. 1.**
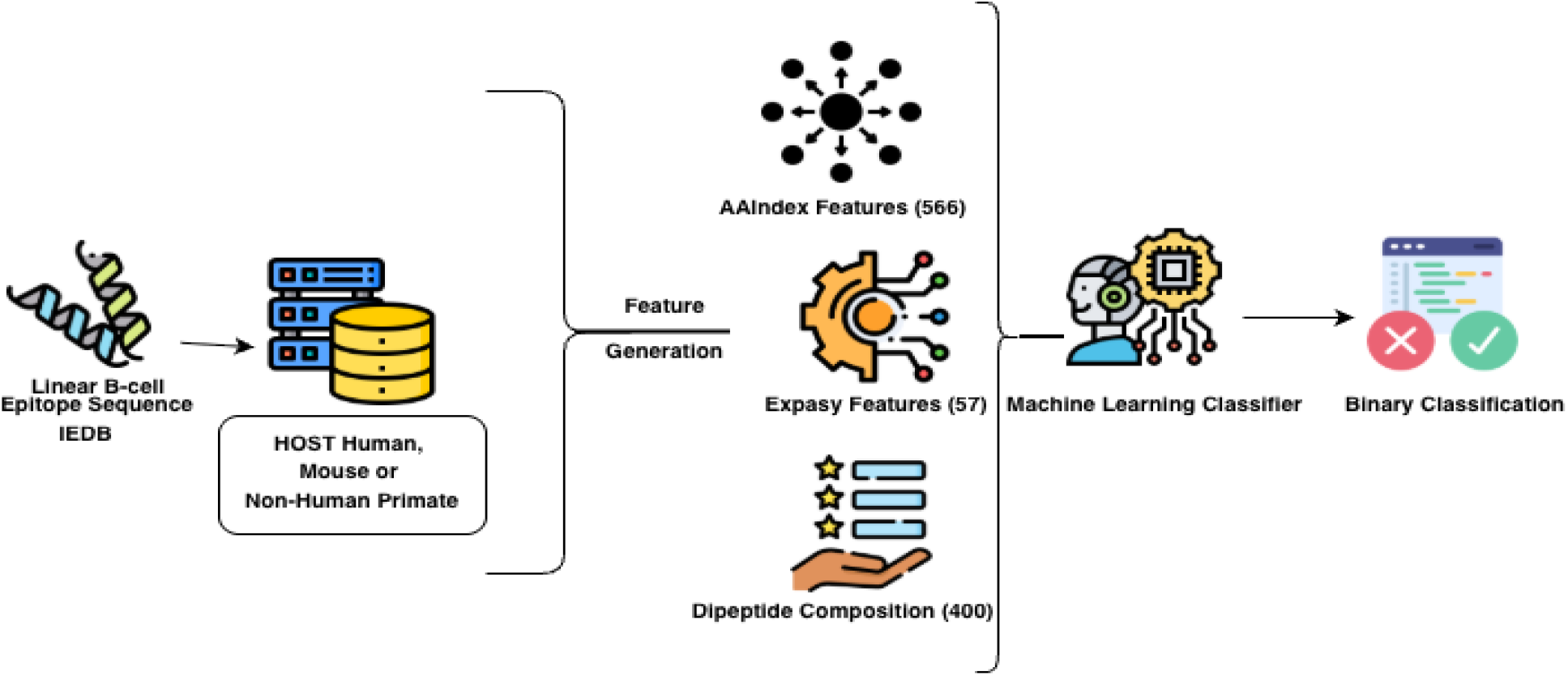
HALPred-B (host-aware linear B-cell epitope prediction) pipeline, where IEDB data is partitioned by host, preprocessed, and encoded into a 1023-dimension feature vector. All five machine learning classifiers are trained and evaluated separately per host.

### A. Dataset Curation from IEDB

All data were obtained from the Immune Epitope Database (IEDB) [24], querying exclusively for linear B-cell assay entries with binary positive or negative outcomes. After applying host-based filters, the final dataset comprises:

- **Human host:** 173,273 positive (epitope) and 222,012 negative (non-epitope) sequences.
- **Mouse host:** 13,909 positives and 13,875 negatives.
- **Non-human primate (NHP) host:** 4,484 positives and 5,991 negatives.

The class imbalance in the human dataset (ratio ≈ 1:1.28) is mild but non-trivial; mouse and NHP datasets are near-balanced by construction. Maintaining host separation throughout the pipeline is essential to prevent immunological context from leaking across organism boundaries.

### B. Data Preprocessing

Raw IEDB sequences undergo four preprocessing steps. First, sequences containing ambiguous amino acid codes (B, J, O, U, X, Z) or non-standard characters are discarded. Second, exact duplicates and entries with contradictory labels across different assays are removed. Third, all accepted sequences are length-normalized to a fixed window of *L* = 20 amino acids by truncating longer peptides from the C-terminus and zero-padding shorter ones; this uniform length is required for the fixed-dimensional AAIndex and dipeptide feature encodings. Fourth, sequence labels are validated to ensure that only entries consistently classified as positive or negative across independent experimental replications are retained, improving label reliability.

### C. Host-Stratified Dataset Partitioning

Each host-specific dataset is split independently into training (80%) and test (20%) subsets using stratified random sampling to preserve class proportions. No data are shared across host partitions at any stage—training, validation, or testing. To address residual class imbalance in the training portion, we apply the Synthetic Minority Over-sampling Technique (SMOTE) [25] with *k* = 5 nearest neighbors to the training set only. The held-out test sets are left unmodified to ensure that evaluation reflects true deployment conditions.

### D. Feature Extraction

Each preprocessed peptide is encoded into a 1023-dimensional feature vector by concatenating three complementary representations:

1. **AAIndex physicochemical descriptors (566 dimensions)**. For each sequence position, residue-level scores from AAIndex1 [26] are aggregated (mean-pooled) across the sequence to obtain a 566-dimensional vector encoding hydrophobicity, charge, flexibility, and surface accessibility, among other biophysical properties.
2. **ExPASy biochemical descriptors (57 dimensions)**. Global sequence-level properties computed via Prot-Param—including molecular weight, isoelectric point, instability index, aliphatic index, and grand average of hydropathicity (GRAVY)—are concatenated to form a 57-dimensional biochemical summary.
3. **Dipeptide composition (400 dimensions)**. The normalized frequency of all 20 × 20 = 400 possible amino acid dipairs is computed, capturing short-range sequential dependencies and amino acid co-occurrence patterns.

All feature vectors are standardized to zero mean and unit variance using parameters estimated solely from the respective host’s training partition to prevent data leakage.

### E. Classifier Training (Per Host)

Five standard classifiers are trained independently for each host:

- **Random Forest (RF):** 500 estimators, maximum depth unconstrained, Gini impurity criterion.
- **Support Vector Machine (SVM):** Radial basis function (RBF) kernel, *C* = 1.0, *γ* = scale.
- **Gradient Boosting (GBC):** 200 estimators, learning rate 0.1, maximum depth 5.
- **XGBoost:** 300 estimators, learning rate 0.05, maximum depth 6, subsample 0.8.
- **K-Nearest Neighbors (KNN):** *k* = 7, Euclidean distance, uniform weights.

All hyperparameters were selected via 5-fold stratified cross-validation on the respective host’s training set, with AUC-ROC as the optimization criterion.

### F. Host-Aware Feature Reweighting (HAFR)

Training a classifier on a host-specific dataset yields an implicit measure of feature importance. HAFR exploits this to construct a refined feature representation that is better calibrated to the host’s immunogenic signals. The procedure operates in three steps, formally stated in Algorithm 1.

Let **x** ∈ ℝ^*d*^ a feature vector (*d* = 1023) and let *f*_*h*_ be a tree-based classifier (RF, GBC, or XGBoost) trained on host *h*. Feature importance scores ***ϕ*** = (*ϕ*_1_, …, *ϕ*_*d*_) are extracted from *f*_*h*_ using the mean decrease in impurity. These are normalized to a probability simplex:

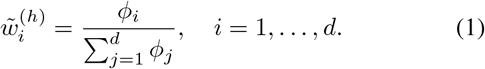

The refined feature vector for any peptide is then:

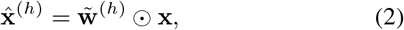

where ⊙ denotes element-wise multiplication. Classifiers are then retrained on these re-weighted feature vectors. For SVM and KNN—which do not natively provide impurity-based importance—feature weights derived from the best tree ensemble on the same host (XGBoost) are used as a proxy.

HAFR is host-specific by construction: the weight vector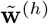 is computed separately for each host and encodes which biochemical and compositional signals are most predictive in that organism’s immunological context. It does not add new features; rather, it suppresses host-irrelevant dimensions and amplifies host-informative ones, leading to a more discriminative input space for all downstream classifiers.

#### Algorithm 1

Host-Aware Feature Reweighting (HAFR)

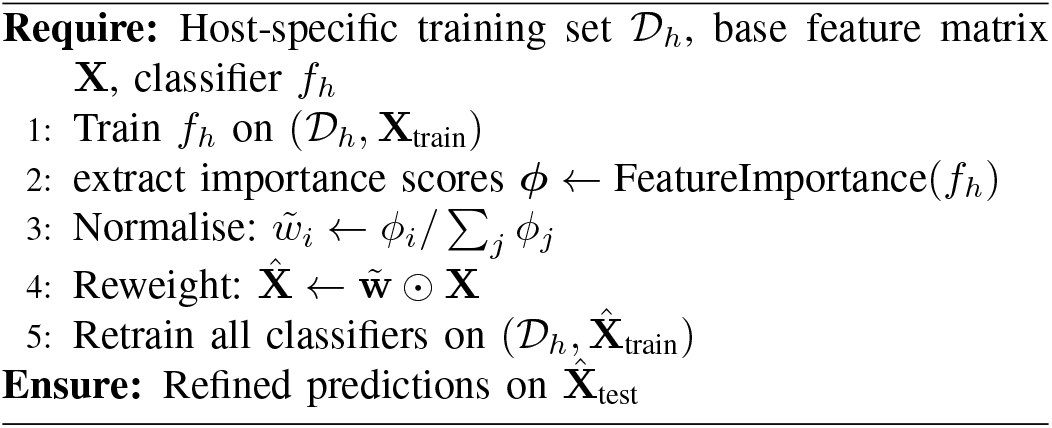

### G. Evaluation Metrics

Models are evaluated on the held-out test set of the corresponding host. Reported metrics include accuracy, precision, recall, F1-score, Matthews Correlation Coefficient (MCC) [27], and AUC-ROC. MCC is particularly valuable here because it accounts for class imbalance in a single scalar measure. All results are reported as the mean over five independent runs with different random seeds to assess stability.

## V. Results and Analysis

### A. Cross-Host Performance Comparison (Baseline)

Table I shows the accuracy of all five classifiers on the three host datasets before HAFR application. A clear trend can be seen with the best accuracy on the human dataset and a decrease in accuracy on the mouse and NHP datasets. For human data, accuracy varies from 79.80% using SVM to 86.07% using XGBoost. In contrast, the mouse achieves only 66.65-69.17% accuracy, some 15-17% points lower than the best human. NHP performs slightly better than mouse, around 69–72%, possibly because its dataset size and sequence diversity are in between the human and mouse datasets. Interestingly, the best accuracy on human data is given by XGBoost, while it performs worst on mouse data, which suggests overfitting to human-specific sequence patterns. This gap is shown in the form of a heatmap in Figure 2. The lower intensity for mouse and NHP further supports that the performance drop is due to the host and not a single classifier.

**TABLE I:** Accuracy comparison of ml classifiers across different host organisms (baseline, without hafr).

| Classifier | Human Host | Mouse Host | Non-Human Primate Host |
| --- | --- | --- | --- |
| Random Forest (RF) | 84.82% | 67.73% | 71.31% |
| Support Vector Machine (SVM) | 79.80% | 68.78% | 69.36% |
| Gradient Boosting (GBC) | 85.79% | **69.17%** | 70.60% |
| XGBoost | **86.07%** | 66.65% | **71.65%** |
| K-Nearest Neighbours (KNN) | 83.45% | 68.45% | 70.22% |

**Fig. 2.**
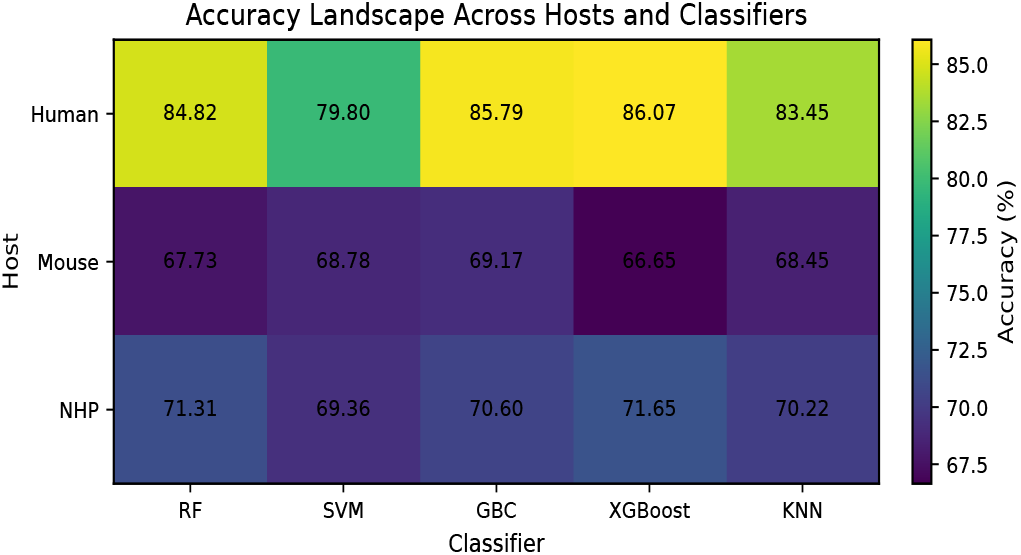
Heatmap of classifier accuracy across host organisms.

### B. Best-Performing Classifier: XGBoost on Human Host

Table II presents the full evaluation of XGBoost on the human host test set. The AUC-ROC of 0.93 shown in Figure 3 indicates strong discriminative ability, and the balanced precision (0.86) and recall (0.85) confirm that performance is not inflated by class-majority bias. The MCC of 0.5441 further validates this: an MCC above 0.5 is considered substantially informative for binary biomedical classification tasks [26].

**TABLE II:** Full evaluation of xgboost on the human host test set.

| Metric | Value |
| --- | --- |
| Accuracy | **86.07%** |
| Precision | 0.86 |
| Recall | 0.85 |
| F1-Score | 0.86 |
| AUC-ROC | 0.93 |
| MCC | 0.5441 |

**Fig. 3.**
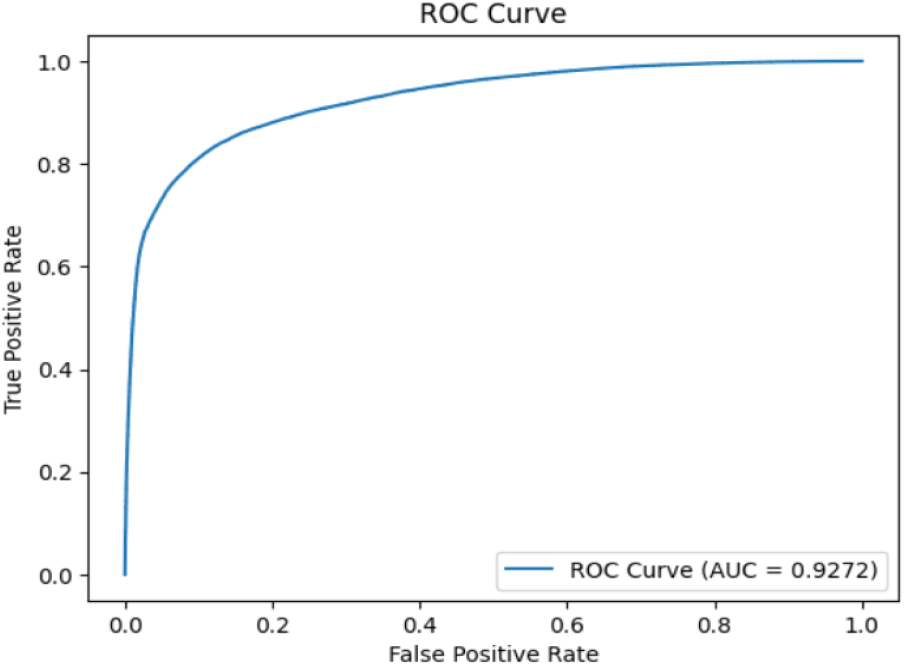
AUC-ROC curve for XGBoost on the human host test set (AUC = 0.93).

### C. Effect of Host-Aware Feature Reweighting (HAFR)

Table III shows a comparison of baseline accuracy versus HAFR-augmented accuracy for all classifier-host combinations. HAFR consistently improves with accuracy gains between 2.4 and 3.6%. The improvements are largest on the NHP dataset, where the lack of data makes the baseline feature space noisier and leaves more room for the amplification of host-specific signals. On the human dataset, which already has a high baseline, HAFR adds 2.4-3.4% showing that even a well-sampled human model benefits from down-weighting immunologically less relevant dimensions. Importantly, HAFR does not fully close the cross-host gap. The top NHP accuracy (75.25% with XGBoost) is still more than 14 points below the top human accuracy after reweighting. This remaining gap is due to fundamental limitations that HAFR cannot overcome: the lack of explicit host-specific biological features (B-cell receptor diversity, post-translational environments, antigen processing machinery) and the relatively small size of the mouse and NHP training sets.

**TABLE III:** Accuracy before and after applying hafr across all classifiers and host organisms.

| Classifier | Human Host |  | Mouse Host |  | Non-Human Primate Host |  |
| --- | --- | --- | --- | --- | --- | --- |
|  | Baseline | +HAFR | Baseline | +HAFR | Baseline | +HAFR |
| Random Forest (RF) | 84.82% | 88.18% | 67.73% | 70.69% | 71.31% | 74.33% |
| SVM | 79.80% | 82.21% | 68.78% | 71.34% | 69.36% | 72.32% |
| Gradient Boosting (GBC) | 85.79% | 89.05% | 69.17% | 72.65% | 70.60% | 73.90% |
| XGBoost | 86.07% | 89.27% | 66.65% | 69.91% | 71.65% | 75.25% |
| K-Nearest Neighbours (KNN) | 83.45% | 86.07% | 68.45% | 70.02% | 70.22% | 73.14% |

### D. Discussion of Host-Dependent Performance

This trend in performance can be explained by three related factors.

#### Dataset size and sampling diversity

The human dataset is much larger than the NHP dataset, which provides the model with more extensive experience with antigenic sequence patterns. More examples will help classifiers to learn more stable decision boundaries even if the underlying features are not perfect.

#### Sequence distribution shift

The epitope sequences are not the same in all hosts. They have different amino acid compositions and dipeptide patterns, in part because different hosts have different pathogen histories. Thus, a model trained mainly on human epitopes may observe different sequence distribution when tested on mouse or NHP data, which leads to lower confidence and lower accuracy.

#### Not enough host-aware features

AAIndex, ExPASy and dipeptide features characterize peptide chemistry, but not host-specific immune biology. They lack factors such as antibody CDR-loop diversity, somatic hypermutation, and antigen-processing conditions like endosomal pH and protease activity. So HAFR can retrieve some signal, but it is not able to fully address the lack of host-biological information.

## VI. Conclusion

In this study, we present HALPred-B, a host-stratified evaluation framework for linear B-cell epitope prediction, and demonstrate that host organism can have a significant effect on model performance. We evaluated five machine learning classifiers using independently curated human, mouse, and non-human primate (NHP) datasets from IEDB and found that regardless of the classifier chosen, the prediction accuracy on non-human hosts remained 15–20 percentage points lower than the best human-host result. This gap suggests that current limitations are not only algorithmic but also biological and data-driven, arising from unequal host-specific dataset sizes, cross-host sequence distribution shifts, and the absence of explicit host-biological information in standard feature representations. To tackle this problem, we proposed HAFR, a post-training feature reweighting strategy that amplifies host-informative feature dimensions. HAFR consistently gained 2–4 percentage points in accuracy, suggesting the existence of host-related signals, but they are weakly captured by generic sequence features. However, these modest gains also show that reweighting alone is not enough.

Multi-task or meta-learning architectures with shared cross-host representations and host-specific prediction heads may further improve data efficiency for underrepresented hosts, while protein language model embeddings such as ESM-2 may help to capture latent evolutionary and host-relevant sequence patterns. Overall, our results question the universal-model assumption in epitope prediction and set host-aware evaluation as a necessary benchmark. Balanced, publicly available host-stratified datasets, particularly for NHP and other underrepresented organisms, will be critical for rigorous and biologically meaningful assessment.

## References

[1] L. Potocnakova, M. Bhide, and L. B. Pulzova, “An introduction to B-cell epitope mapping and in silico epitope prediction,” J. Immunol. Res., vol. 2016, p. 6760830, 2016.

[2] S. K. Dhanda, S. S. Usmani, P. Agrawal, G. Nagpal, A. Gautam, and G. P. Raghava, “Novel in silico tools for designing peptide-based subunit vaccines and immunotherapeutics,” Brief. Bioinformatics, vol. 18, no. 3, pp. 467–478, 2017.

[3] E. A. Emini, J. V. Hughes, D. S. Perlow, and J. Boger, “Induction of hepatitis A virus-neutralizing antibody by a virus-specific synthetic peptide,” J. Virol., vol. 55, no. 3, pp. 836–839, 1985.

[4] R. Vita et al., “The Immune Epitope Database (IEDB) 3.0,” Nucleic Acids Res., vol. 43, no. D1, pp. D405–D412, 2015.

[5] J. M. R. Parker, D. Guo, and R. S. Hodges, “New hydrophilicity scale derived from high-performance liquid chromatography peptide retention data: correlation of predicted surface residues with antigenicity and X-ray-derived accessible sites,” Biochemistry, vol. 25, no. 19, pp. 5425–5432, 1986.

[6] P. A. Karplus and G. E. Schulz, “Prediction of chain flexibility in proteins: a tool for the selection of peptide antigens,” Naturwissenschaften, vol. 72, no. 4, pp. 212–213, 1985.

[7] A. S. Kolaskar and P. C. Tongaonkar, “A semi-empirical method for prediction of antigenic determinants on protein antigens,” FEBS Lett., vol. 276, no. 1–2, pp. 172–174, 1990.

[8] T. P. Hopp and K. R. Woods, “Prediction of protein antigenic determinants from amino acid sequences,” Proc. Natl. Acad. Sci. USA, vol. 78, no. 6, pp. 3824–3828, 1981.

[9] C. P. Toseland et al., “AntiJen: a quantitative immunology database integrating functional, thermodynamic, kinetic, biophysical, and cellular data,” Immunome Res., vol. 1, no. 1, p. 4, 2005.

[10] Y. EL-Manzalawy, D. Dobbs, and V. Honavar, “Predicting linear B-cell epitopes using string kernels,” J. Mol. Recognit., vol. 21, no. 4, pp. 243–255, 2008.

[11] J. E. P. Larsen, O. Lund, and M. Nielsen, “Improved method for predicting linear B-cell epitopes,” Immunome Res., vol. 2, no. 1, p. 2, 2006.

[12] H. Singh, H. R. Ansari, and G. P. S. Raghava, “Improved method for linear B-cell epitope prediction using antigen’s primary sequence,” PLOS ONE, vol. 8, no. 5, p. e62216, 2013.

[13] M. J. Blythe and D. R. Flower, “Benchmarking B cell epitope prediction: underperformance of existing methods,” Protein Sci., vol. 14, no. 1, pp. 246–248, 2005.

[14] S. Saha and G. P. S. Raghava, “Prediction of continuous B-cell epitopes in an antigen using recurrent neural network,” Proteins: Struct. Funct. Bioinformatics, vol. 65, no. 1, pp. 40–48, 2006.

[15] B. Yao, L. Zhang, S. Liang, and C. Zhang, “SVMTriP: a method to predict antigenic epitopes using support vector machine to integrate tripeptide similarity and propensity,” PLOS ONE, vol. 7, no. 9, p. e45152, 2012.

[16] B. M. da Silva, D. B. Ascher, and D. E. V. Pires, “epitope1D: accurate taxonomy-aware B-cell linear epitope prediction,” Brief. Bioinformatics, vol. 24, no. 3, p. bbad114, 2023.

[17] F. Campelo et al., “Phylogeny-aware linear B-cell epitope predictor detects targets associated with immune response to orthopoxviruses,” Brief. Bioinformatics, vol. 25, no. 6, p. bbae527, 2024.

[18] J. Ashford, J. Reis-Cunha, I. Lobo, F. Lobo, and F. Campelo, “Organism-specific training improves performance of linear B-cell epitope prediction,” Bioinformatics, vol. 37, no. 24, pp. 4826–4834, 2021.

[19] T. C. Shih, L. P. Ho, H. Y. Chou, J. L. Wu, and T. W. Pai, “Comprehensive linear epitope prediction system for host specificity in Nodaviridae,” Viruses, vol. 14, no. 7, p. 1357, 2022.

[20] A. Ras-Carmona, H. F. Pelaez-Prestel, E. M. Lafuente, and P. A. Reche, “BCEPS: a web server to predict linear B cell epitopes with enhanced immunogenicity and cross-reactivity,” Cells, vol. 10, no. 10, p. 2744, 2021.

[21] K. Murphy and C. Weaver, Janeway’s Immunobiology, 9th ed. New York: Garland Science, 2016.

[22] A. K. Abbas, A. H. Lichtman, and S. Pillai, Cellular and Molecular Immunology, 9th ed. Philadelphia: Elsevier, 2018.

[23] W. E. Paul, Fundamental Immunology, 7th ed. Philadelphia: Lippincott Williams & Wilkins, 2012.

[24] R. Vita et al., “The Immune Epitope Database (IEDB) 3.0,” Nucleic Acids Res., vol. 43, no. D1, pp. D405–D412, 2015.

[25] N. V. Chawla, K. W. Bowyer, L. O. Hall, and W. P. Kegelmeyer, “SMOTE: synthetic minority over-sampling technique,” J. Artif. Intell. Res., vol. 16, pp. 321–357, 2002.

[26] P. Gautam and P. Mitra, ‘“ConfPred: ML-based conformational B-cell epitope prediction using novel features,” in Proceedings of the 12th International Conference on Bioinformatics Research and Applications, pp. 103–110, 2025.

[27] D. Chicco and G. Jurman, “The advantages of the Matthews correlation coefficient (MCC) over F1 score and accuracy in binary classification evaluation,” BMC Genomics, vol. 21, no. 1, p. 6, 2020.

